## Supplementary Figures for "Post-transcriptional splicing can occur in a slow-moving zone around the gene"

### Supplementary Figure 1

**A**

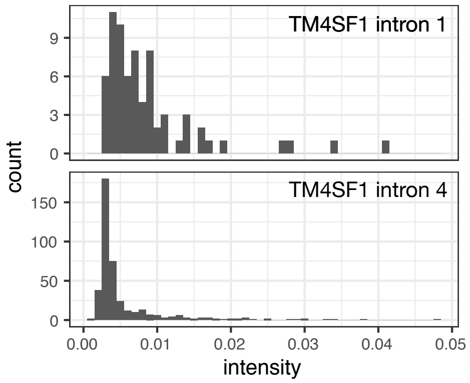


**Supplementary Figure 1: Transcription site choice and defining post-transcriptionality.**
A. Histograms of intron intensities for TM4SF1 intron 1 and intron 4, before, after, and during defining a global thresholding cutoff and other transcription site selection methods, as well as dispersal graphs generated based on those transcription site selections.
B. Number of dispersed pre-mRNA per cell.
C. Parameter sweep of colocalization threshold and associated dispersal graphs.
D. Example classification of pre-mRNA as either transcription sites or dispersed pre-mRNA.
E. Distances between 5’ and 3’ ends of RNA as detected by RNA FISH for either the 3’ or 5’ end of the RNA of interest.
F. Example classification of RNA as “distal” or “proximal” to the transcription site
G. Histogram showing type of pre-mRNA as defined by RNA FISH for stated introns in FKBP5, as a function of distance to the nearest transcription site.

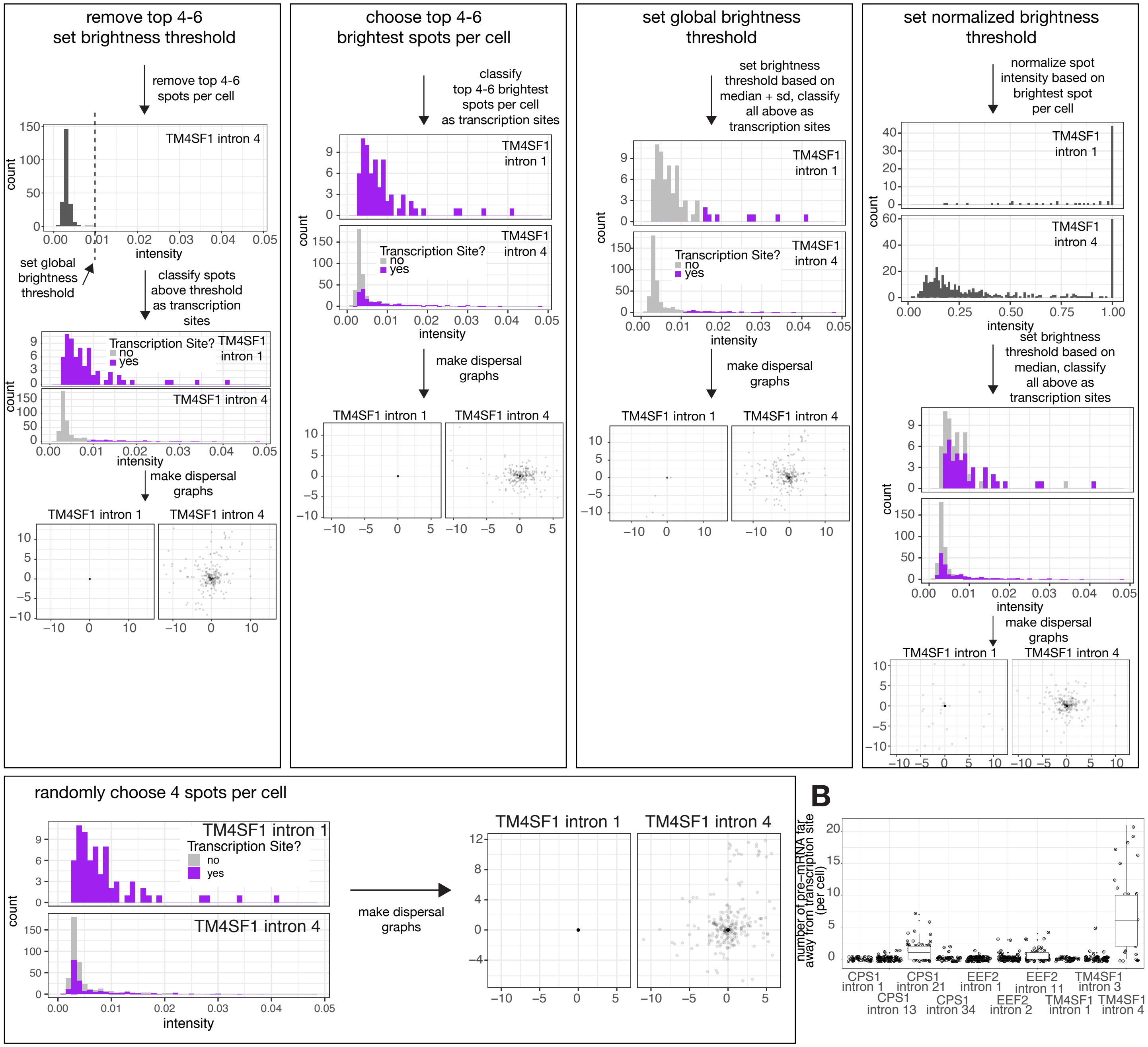

### Supplementary Figure 1

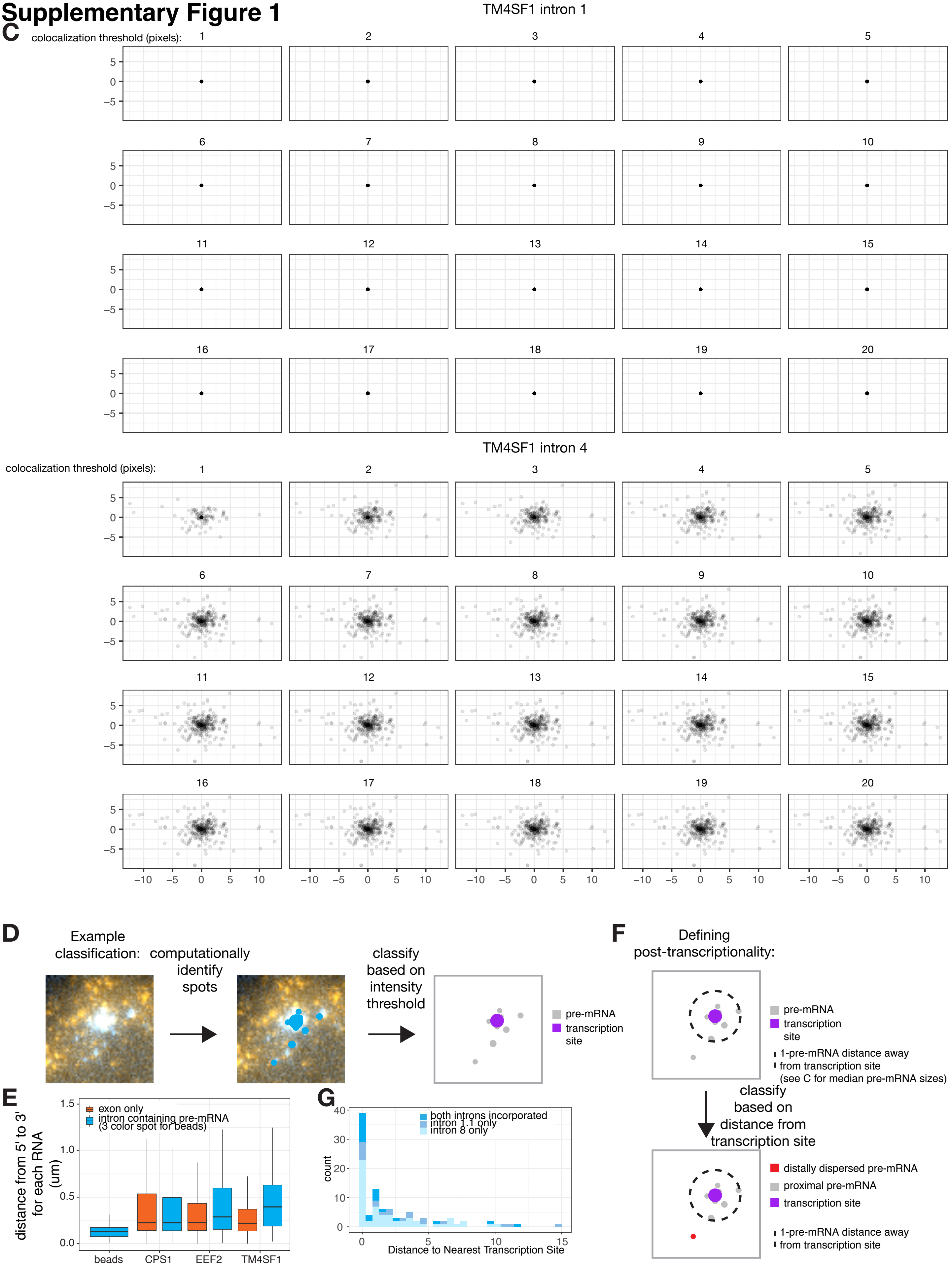

### Supplementary Figure 2

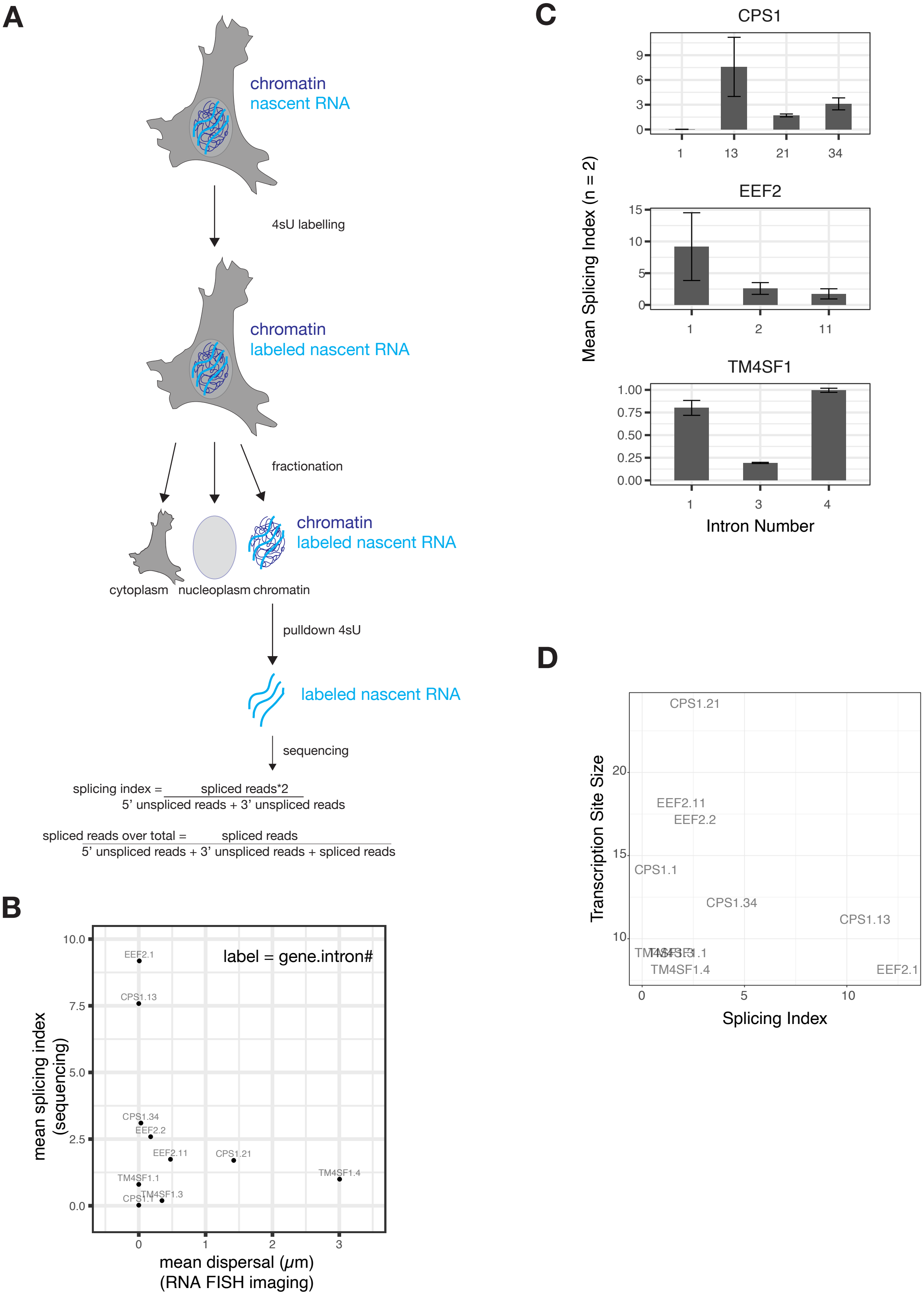

### Supplementary Figure 3

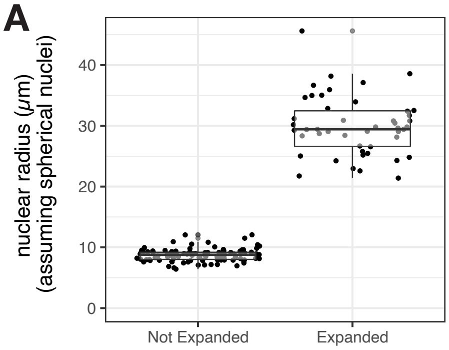


**Supplementary Figure 3: Expansion microscopy yields a 4.65 fold linear expansion and expands isotropically.** A. Compari- son of radii of nuclei (based on DAPI staining, and assuming spherical nuclei) before and after expansion. B. Images of the same cell before and after expansion, with or without pladeinolide B treatment (as noted). Scale bars = 5μm. C. Transcription site area (microns squared) for 5' and 3' probes, with and without dye swap.

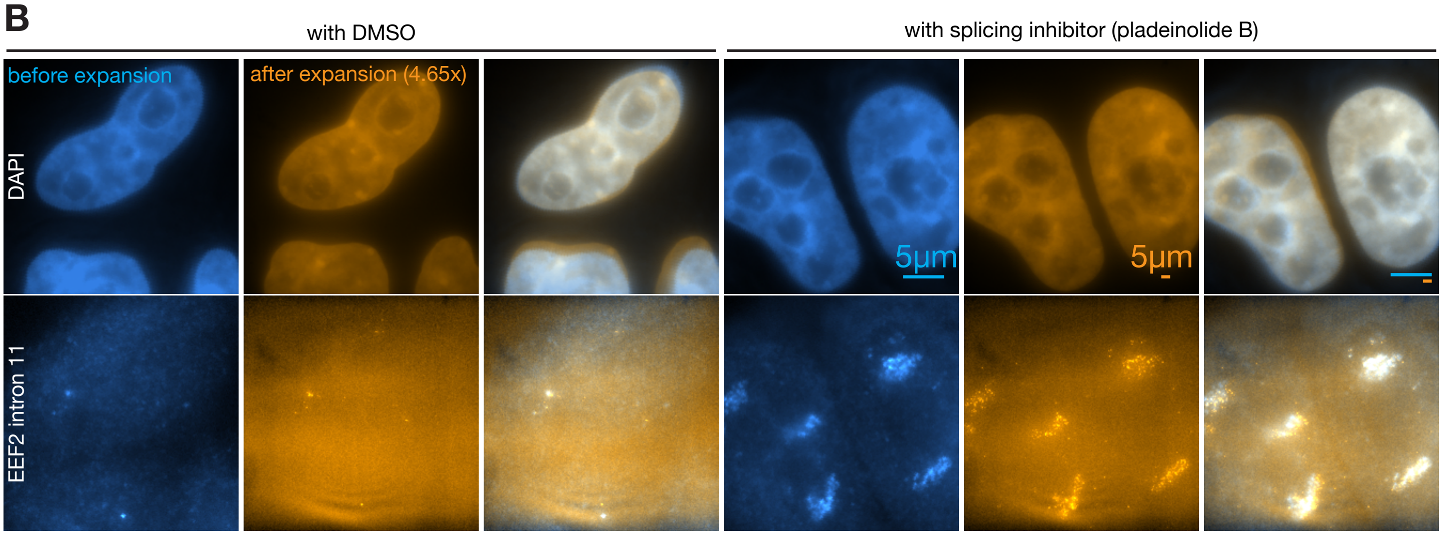


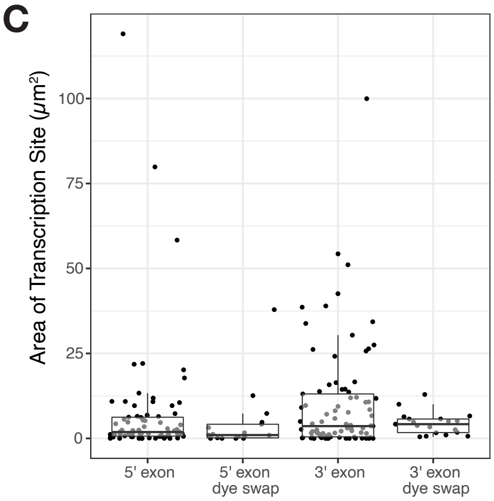

### Supplemental Figure 4

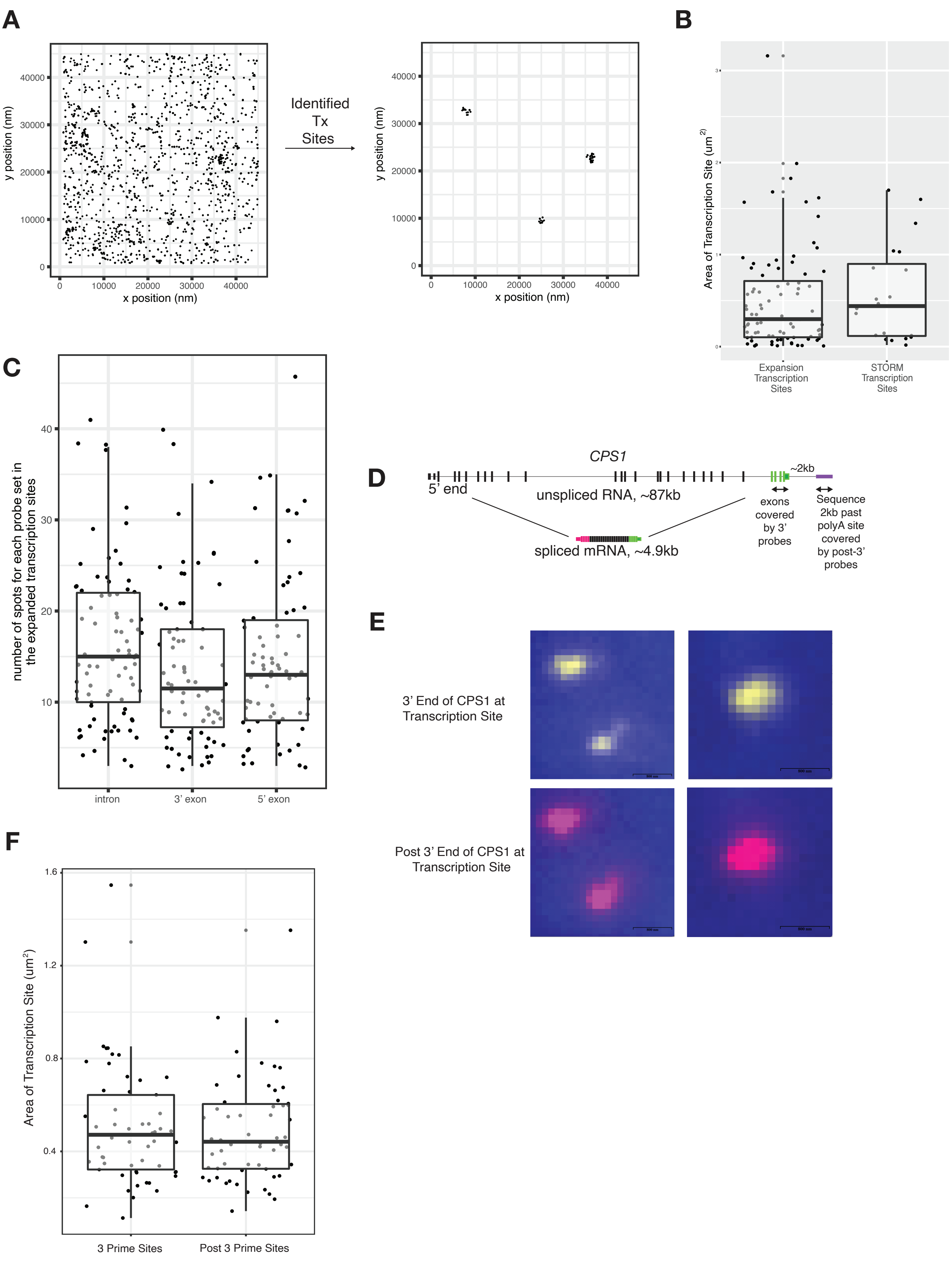

### Supplementary Figure 5

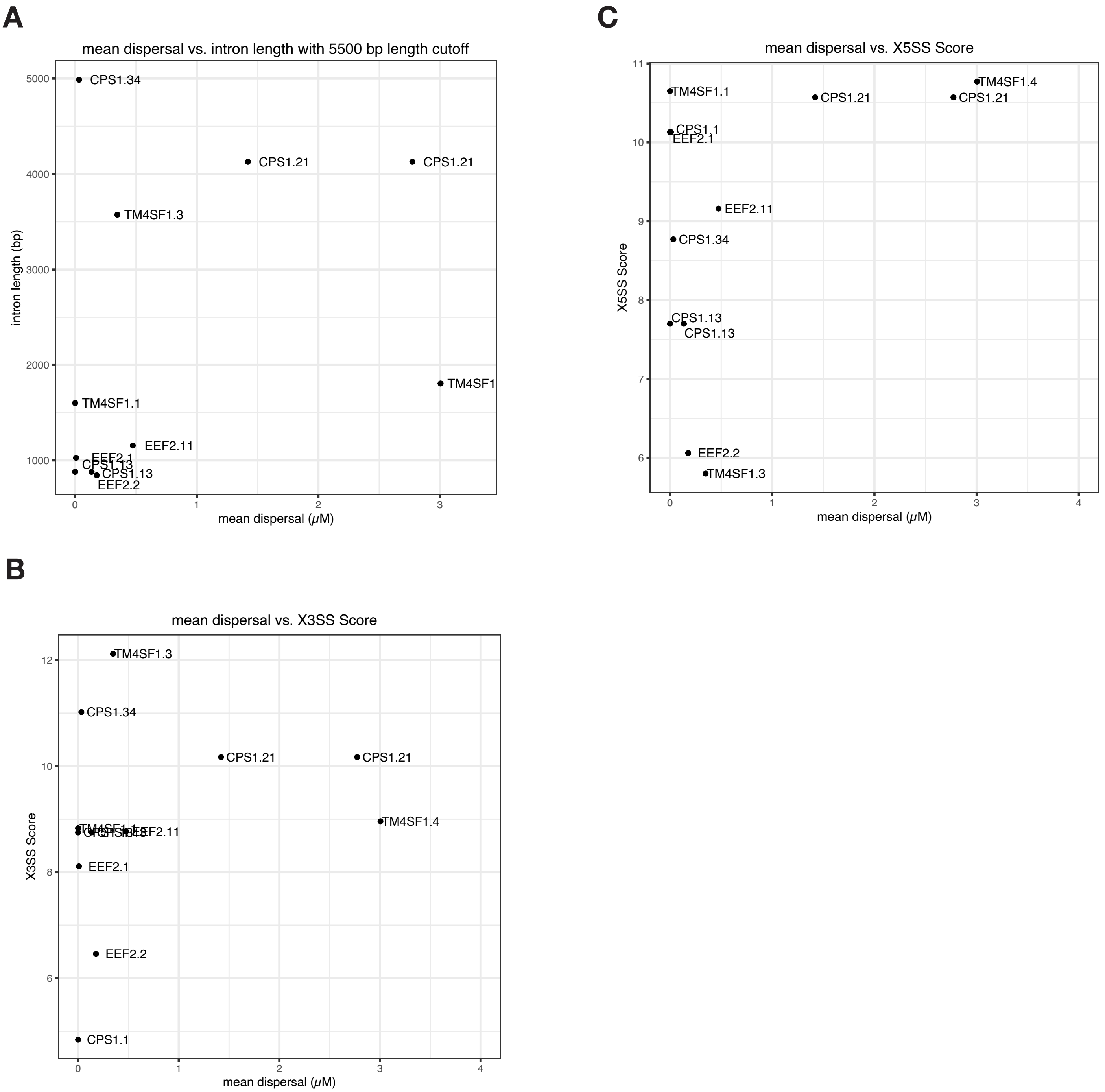

#### Supplementary Figure 6

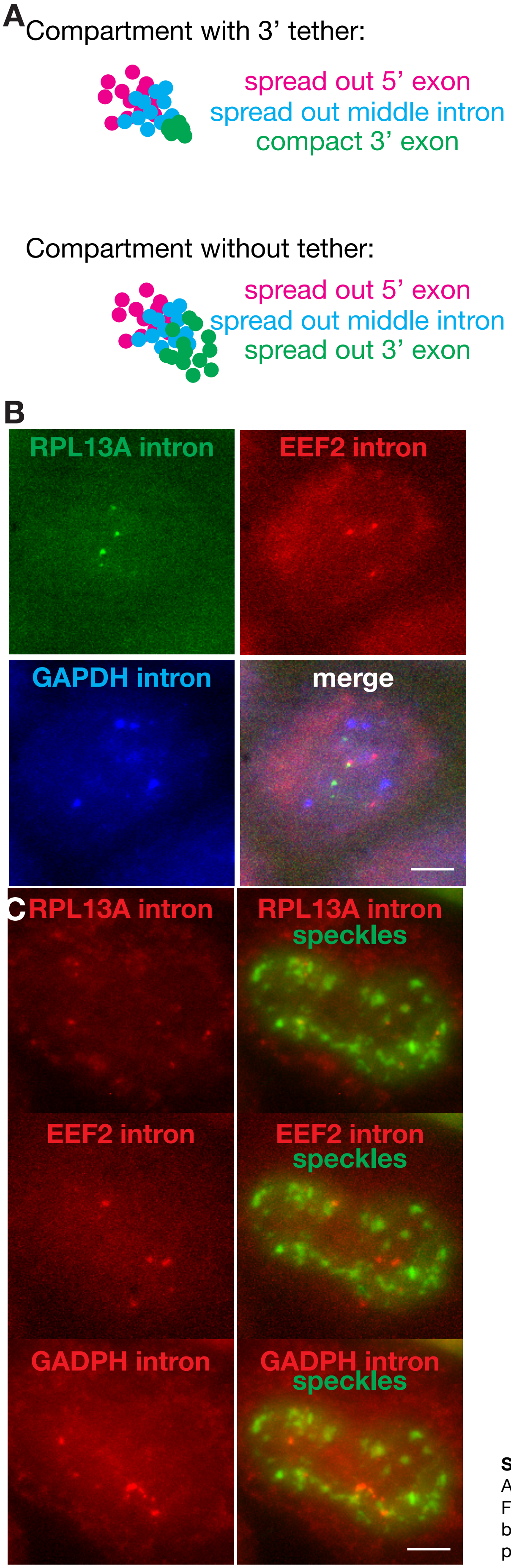


**Supplementary Figure 6: Compartmentalization genes before splicing inhibition.** A. Schematic of compartmentalization phenotype with and without tether. B. RNA FISH of RPL13A, EEF2, and GAPDH introns before pladienolide B treatment. Scale bar = 5 μm. C. Combined RNA FISH for the stated introns and IF for SC35, before pladienolide B treatment. Scale bar = 5 μm.
