## Supplementary figures and images for "Post-transcriptional splicing can occur in a slow-moving zone around the gene"

### Supplemental Movie 1

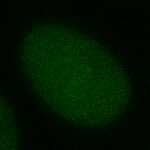

### Supplemental Movie 2

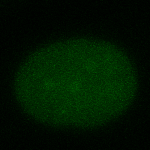

### Supplemental Movie 3

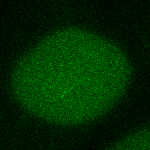
